# Examining go-or-grow using fluorescent cell-cycle indicators and cell cycle-inhibiting drugs

**DOI:** 10.1101/797142

**Authors:** Sean T. Vittadello, Scott W. McCue, Gency Gunasingh, Nikolas K. Haass, Matthew J. Simpson

## Abstract

The go-or-grow hypothesis states that adherent cells undergo reversible phenotype switching between migratory and proliferative states, with cells in the migratory state being more motile than cells in the proliferative state. Here we examine go-or-grow in 2-D *in vitro* assays using melanoma cells with fluorescent cell-cycle indicators and cell cycle-inhibiting drugs. We analyse the experimental data using single-cell tracking to calculate mean diffusivities, and compare motility between cells in different cell-cycle phases and in cell-cycle arrest. Unequivocally, our analysis does not support the go-or-grow hypothesis. We present clear evidence that cell motility is independent of the cell-cycle phase, and non-proliferative arrested cells have the same motility as cycling cells.

---

The *go-or-grow* hypothesis, also referred to as the *migration/proliferation dichotomy* or the *phenotype switching model*, proposes that adherent cells reversibly switch between migratory and proliferative phenotypes [1], exhibiting higher motility in the migratory state as motile cells are not using free energy for proliferation [1–5]. Previous experimental investigations of the go-or-grow hypothesis are conflicting, as some studies support the hypothesis [1, 6, 7] while others refute it [8–10].

Go-or-grow was initially proposed as an explanation for the apparent mutual exclusivity of migration and proliferation for astrocytoma cells, first in 2-D *in vitro* experiments [7], and later for *in vivo* investigations [6]. In these early studies, claims for evidence of go-or-grow are based on the comparison of the subpopulation of cells at the perimeter of the cell population, where cells are considered to be invasive, with the subpopulation of cells in the central region, where cells are considered non-invasive. Data suggest that the proliferation rate is lower at the perimeter and higher in the centre, leading to the assertion that the more migratory cells are less proliferative. The experimental data, however, only indicate that the subpopulation at the perimeter is less proliferative as a whole compared with the centre, and therefore we cannot conclude definitively that the more migratory cells are less proliferative.

To test for evidence of go-or-grow it is necessary to look at the single-cell level, as in subsequent studies [8–10], where single-cell tracking is used with single-cell migration measured in terms of the net displacement of the cell trajectory. These three studies, none of which support go-or-grow, involve 2-D and 3-D *in vitro* experiments with medulloblastoma cells [10], 2-D *in vitro* experiments with mesothelioma, melanoma, and lung cancer cells [9], and 2-D and 3-D *in vitro* experiments with melanoma cells [8]. Studies of tumour heterogeneity in melanoma suggest that cells may reversibly switch between invasive and proliferative phenotypes [1]. As melanoma is highly metastatic, forms tumours that are very heterogeneous, and is well known to respond to MAPK inhibitors which induce G1 arrest [11,12], melanoma cells are a prime candidate for studying the go-or-grow hypothesis.

Confirmation of go-or-grow would have important implications for anti-cancer treatments employing cell cycle-inhibiting drugs. For most eukaryotic cells, the cell cycle is a sequence of four discrete phases (Fig. 1a), namely gap 1 (G1), synthesis (S), gap 2 (G2) and mitosis (M). Cell-cycle arrest (Fig. 1d), which occurs when progression through the cell cycle halts [13], can be induced by cell cycle-inhibiting drugs [8, 14, 15]. An arrested cell is not proliferative, so the cell’s free energy could be utilised for migration, potentially leading to an exacerbation of metastasis [3].

**Figure 1:**
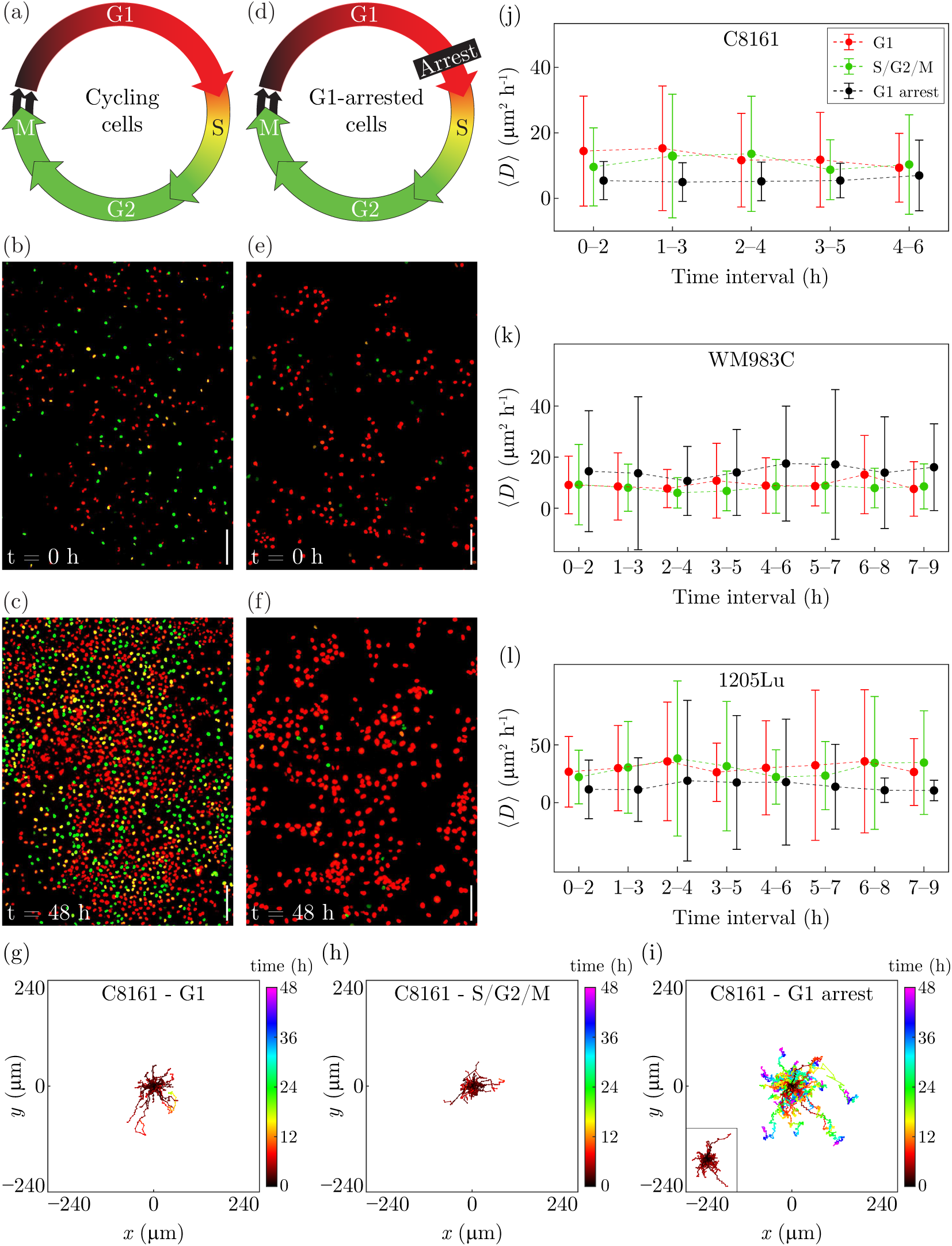
Experimental data and mean diffusivities. (a) The cell cycle, indicating the colour of FUCCI in each phase. (b)–(c) Experimental images of cycling C8161 cells; cell counts at 0 and 48 h are 331 and 1878, respectively. (d) The cell cycle, indicating the colour of FUCCI in each phase together with arrest in G1. (e)–(f) Experimental images of C8161 cells in G1 arrest (30 nM trametinib); cell counts at 0 and 48 h are 261 and 469, respectively. (g)–(i) 50 cell trajectories of G1 cycling, S/G2/M cycling and G1-arrested (30 nM trametinib) C8161 cells, respectively. (j)–(l) No difference in mean diffusivity, ⟨*D*⟩, for C8161, WM983C and 1205Lu cells, respectively. For each 2-h time interval, ⟨*D*⟩ is the mean of all individual diffusivities *D* corresponding to cells with trajectories within the time interval. In each case we show ⟨*D*⟩, and report the variability using ⟨*D*⟩ plus or minus the sample standard deviation. Data for each experimental condition are offset with respect to the time-interval axis for clarity. Scale bar 200 *µ*m.

The go-or-grow hypothesis also has important implications for mathematical models of collective cell invasion in a population of migratory and proliferative cells. Such models of cell invasion are often based on the Fisher–Kolmogorov–Petrovskii–Piskunov (FKPP) equation [16–19],

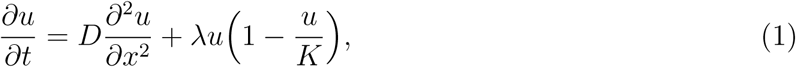

where *x* is position, *t* is time, *u*(*x, t*) > 0 is cell density, *D* > 0 is the diffusivity, *λ* > 0 is the proliferation rate, and *K* > 0 is the carrying-capacity density. Eq. (1) and related adaptations, including stochastic analogues [20, 21], have been successfully used to model cell migration *in vitro* and *in vivo* [22–26]. A key assumption underlying these models is that *D* is independent of the cell-cycle phase, which may not hold if cells are subject to go-or-grow as then a cycling, therefore non-arrested, cell may become less motile as it progresses through the cell cycle and nears cell division [8].

In this work we rigorously examine the go-or-grow hypothesis for adherent melanoma cells, for which phenotype switching between migratory and proliferative states is proposed to occur [1]. We use melanoma cell lines in this study as melanoma is the *prototype* for the phenotype switching model and is highly responsive to G1 arrest-inducing MEK inhibitors, such as trametinib. Melanoma cells are therefore an ideal candidate for studying go-or-grow [1,3,27]. Our experimental data are obtained from single-cell tracking in 2-D *in vitro* assays. We conduct our experiments in 2-D as it is the natural situation in which to commence a new experimental study, before utilising the knowledge gained in more complicated 3-D or *in vivo* experiments. Indeed, experimental studies of cell migration are often conducted in 2-D *in vitro* assays for several reasons: the observed cell migration is partly representative of cell migration *in vivo*; the assays are amenable to standard laboratory techniques, such as live-cell microscopy; and the relative ease of image analysis, such as cell counting and single-cell tracking [28–30]. Further, cell migration in 3-D may be affected by the properties of a 3-D matrix, which is not present in 2-D assays. For example, cell migration in 3-D through constricting pores can damage the nucleus and thereby cause a delay in cell division as the nucleus undergoes repair, which could be interpreted incorrectly as evidence for go-or-grow [31].

We employ fluorescent ubiquitination-based cell cycle indicator (FUCCI) [32], which consists of two reporters enabling visualisation of the cell cycle of individual live cells: when the cell is in G1 the nucleus fluoresces red, and when the cell is in S/G2/M the nucleus fluoresces green (Fig. 1(a)). During early S, called eS, both of the red and green reporters are active producing yellow. FUCCI allows us to study cell motility in G1 separately from cell motility in S/G2/M [8, 22, 33, 34]. Specifically, we investigate cycling cells for differences in motility when the cells are in G1 compared with S/G2/M. Further, given the potential for an arrested cell to become more motile, we use a cell cycle-inhibiting drug to effect G1 arrest, and compare the motility of the arrested cells with cycling cells. Note that FUCCI does not provide delineation of S, G2 and M, so our motility measurements for these phases are combined into S/G2/M.

Our methodology for examining go-or-grow is novel in a number of ways. We induce G1 arrest in cells to determine whether non-proliferative cells have higher motility than cycling cells. We use experimental data to show that our three cell lines have distinctly different cell-cycle durations, ratios of duration in G1 to S/G2/M, and migration characteristics, all of which may affect motility under the go-or-grow hypothesis. Importantly, the data set we generate and analyse is large: for each cell line and experimental condition we randomly sample 50 single-cell trajectories for analysis out of more than 10^3^ trajectories. In total, we analyse 450 carefully-collected trajectories for evidence of go-or-grow. Using these trajectories we carefully estimate diffusivities by first accounting for anisotropy in the cell migration, so that our estimates are based on time frames for which the cells are undergoing free diffusion.

Our data consist of time-series images, acquired every 15 min for 48 h, from 2-D proliferation assays using the melanoma cell lines C8161, WM983C and 1205Lu [8, 22, 35, 36], which have respective mean cell-cycle durations of approximately 21, 23 and 37 h [8]. The cell lines have very different ratios of durations in G1 to S/G2/M (Supplementary Material). Fig. 1(b)–(c) shows images of an assay with cycling C8161 cells at 0 and 48 h, illustrating the red, yellow and green nuclei corresponding to cells in G1, eS and S/G2/M, respectively. For comparison, Fig. 1(e)–(f) shows images of an assay with G1-arrested C8161 cells treated with the cell cycle-inhibiting drug trametinib (30 nM), illustrating that most cells are arrested in G1, appearing red. We use the lowest possible concentration of trametinib to induce G1 arrest for the experiment duration to minimise other effects. Consequently, each cell eventually returns to cycling, illustrated by the small proportion of green cells (Fig. 1(e)–(f)). These few green cells will eventually divide with both daughter cells arresting in G1. We quantitatively confirm the G1 arrest by comparing the cell counts between the experiments with cycling cells and arrested cells. For the cycling cells there is a 5.7-fold increase in the number of cells over 48 h (Fig. 1(b)–(c)), whereas there is only a 1.8-fold increase in the number of arrested cells over 48 h (Fig. 1(e)–(f)). The 1.8-fold increase in the population of G1-arrested cells is expected as we use the lowest possible concentration of trametinib. Consequently, a small subpopulation of cells may not be arrested at the start of the experiment, and cells may recommence cycling during the experiment, producing a small increase in the population.

For each cell line, we employ single-cell tracking to obtain 50 trajectories of cells for each experimental condition: (i) G1 cycling; (ii) S/G2/M cycling; and (iii) G1 arrest. Each trajectory is selected randomly without replacement from the set of all trajectories for a given cell line and experimental condition. For the cycling cells, trajectories are recorded for the complete duration of the G1 or S/G2/M phase. For the G1-arrested cells, the duration of the trajectory corresponds to the maximum duration that the cell is arrested within the 48-h duration of the experiment (Supplementary Material).

In Fig. 1(g)–(i) we visualise the trajectories for cycling C8161 in G1 and S/G2/M, and C8161 in G1-arrest. The trajectories are translated so that their initial positions are at the origin. The trajectories of the G1-arrested cells are generally much longer than those for the cycling cells, as the arrested cells reside in G1 for a much longer duration than cycling cells reside in G1 or S/G2/M. Specifically, the approximate mean duration of cycling C8161 cells in G1 is 5 h, in S/G2/M is 6 h [8], and for cells in G1 arrest during the 48 h of the experiment is 34 h (Supplementary Material). Therefore, to easily compare the trajectories of G1-arrested cells with cycling cells in G1, we show within the inset the truncated trajectories of the G1 arrested cells. The trajectories are truncated to a duration equal to the mean duration of the corresponding trajectories for cycling cells in G1. Based on these data, the migration is isotropic, without any drift, and independent of the cell cycle phase. We now quantify these observations.

For each cell line and experimental condition, we find that the cell migration is isotropic and directional persistence dissipates within a relatively short lag time of 1 h (Supplementary Material). From each individual cell trajectory we estimate *D*, using the mean square displacement as a function of lag time, within 2-h time intervals. The intervals begin at the initial point of the trajectory, *t* = 0 h, with successive intervals offset by 1 h. We always use lag times from 1–2 h to guarantee the absence of persistence (Supplementary Material). We then calculate the mean diffusivity ⟨*D*⟩ for each time interval by averaging our estimates of *D* for those trajectories that extend to the end of that interval.

Fig. 1(j)–(l) shows, for each cell line, ⟨*D*⟩ for successive time intervals. From these data we arrive at clear conclusions (Supplementary Material), none of which are consistent with the go-or-grow hypothesis:

- For each cell line and experimental condition, there is little variation in ⟨*D*⟩ over time, indicating that there is no appreciable change in motility during each cell-cycle phase and during G1 arrest (Supplementary Material).
- For each cell line, there is little variation in ⟨*D*⟩ between cycling cells in G1, cycling cells in S/G2/M, and G1-arrested cells. The lack of variability in ⟨*D*⟩ is remarkable, and clearly demonstrates that cells in G1 are not more motile than cells in S/G2/M, and that G1-arrested cells at no time become more migratory than the cycling cells.
- Even though our three cell lines have very different proliferation and migration characteristics (Supplementary Material), our estimate of ⟨*D*⟩ is remarkably consistent across the three very different cell lines.

In summary, our analysis of cell migration in 2-D assays using three melanoma cell lines does not support the go-or-grow hypothesis. We find that cell motility is independent of the cell-cycle phase, so that the implication from go-or-grow that cells are more motile in G1 than in S/G2/M when they are nearing cell division is not supported by our data. Notably, there is no change in cell motility when we effect drug-induced G1 arrest in the cells, again displaying a lack of support for the go-or-grow hypothesis.

## Author Contributions

All authors designed the research. STV performed the research. All authors contributed analytic tools and analysed the data. STV wrote the manuscript, and all authors approved the final version of the manuscript.

## Acknowledgments

NKH is a Cameron fellow of the Melanoma and Skin Cancer Research Institute, and is supported by the NHMRC (APP1084893). MJS is supported by the ARC (DP170100474).

