## Supplementary Material 1 for "Examining go-or-grow using fluorescent cell-cycle indicators and cell cycle-inhibiting drugs"

*January 2, 2020*

**References** **29**

### 1 Materials and Methods

#### 1.1 Experiments

Our experimental data consist of microscopy time-series images of two-dimensional cell proliferation assays using the three melanoma cell lines C8161, WM983C and 1205Lu [1–4], which have cell cycle durations of approximately 21, 23 and 37 h, respectively [4]. Here we discuss in detail our experimental set-up and analysis of the time-series images.

##### 1.1.1 Cell culture

The human melanoma cell lines C8161 (kindly provided by Mary Hendrix, Chicago, IL, USA), WM983C and 1205Lu (both kindly provided by Meenhard Herlyn, Philadelphia, PA, USA) were genotypically characterised [5–8], grown as described in [9], and authenticated by STR fingerprinting (QIMR Berghofer Medical Research Institute, Herston, Australia).

We maintain the cell cultures to prevent any induced synchronisation from cell cycle arrest in G1 phase. In general, such induced synchronisation can occur through various culture conditions, namely contact inhibition of proliferation at relatively high population densities [10], decreased pH of the growth medium due to the concentration of acidic cell-metabolites such as lactic acid [11], and reduced availability of nutrients such as serum [12]. We prevent induced synchronisation by passaging the cells every three days, and on the day prior to setting up an experiment, to maintain a subconfluent cell density and a fresh growth medium.

##### 1.1.2 Fluorescent ubiquitination-based cell cycle indicator (FUCCI)

To generate stable melanoma cell lines expressing the FUCCI constructs, mKO2-hCdt1 (30-120) and mAG-hGem (1-110) [13] were subcloned into a replication-defective, self-inactivating lentiviral expression vector system as previously described [14]. The lentivirus was produced by co-transfection of human embryonic kidney 293T cells. High-titer viral solutions for mKO2-hCdt1 (30/120) and mAG-hGem (1/110) were prepared and used for co-transduction into the melanoma cell lines, and subclones were generated by single-cell sorting [4, 9, 15].

##### 1.1.3 Proliferation assay: cycling cells

Cells are seeded from subconfluent culture flasks onto a 24-well plate at a density of $10^4$ cells cm$^{-2}$, with 2.5 ml of medium per well, which is 2.5 times the volume of the standard protocol. After incubating the plate for 24 h at 37 °C with 5% $CO_2$, live-cell images are acquired at 15 min intervals over 48 h at six different positions within the well. For each cell line we performed four independent experiments, with one technical replicate in three experiments and two technical replicates in the fourth experiment. Therefore, for each cell line we have 30 sets of time-series images.

Our preliminary experiments used a standard 1 ml of medium in each well, however the cells started to arrest in G1 around 48 h after seeding, which is likely due to decreased pH of the medium from the lactic acid concentration. G1 arrest is visually obvious when viewing a time series of images of FUCCI cells as the cycling cells are either green or become green according to the duration of G1, while the G1-arrested cells remain red. We therefore performed preliminary tests in an attempt to prevent the cells from arresting in G1 during the experiment, and found that this is possible by increasing the volume of medium in each well to the workable maximum of 2.5 ml, given the volume of each well is 3 ml. The result of the increased volume of medium is that the cells do not begin to arrest in G1 until around 72 h following seeding, which provides us with almost 48 h of imaging using cells that have minimal G1 arrest. Our definition of minimal G1 arrest is that there is no visually detectable arrest due to culture conditions, such as low extracellular pH, throughout the time series images except perhaps within the last hour of the experiment.

##### 1.1.4 Proliferation assay: G1-arrested cells

The experimental set up is identical to that for the cycling cells, except that the MEK inhibitor trametinib, which causes G1 cell-cycle arrest, is added at a concentration of 30 nM. Trametinib is added 3.5 h after seeding, to allow the cells to attach to the plate before becoming affected by the drug. In order to minimise other effects of trametinib we employ the minimum concentration of the drug that causes G1 arrest. Preliminary testing found that the optimal concentration for our cell lines is 30 nM. For each cell line, we performed two independent experiments, with two technical replicates in each experiment. Therefore, for each cell line we have 24 sets of time-series images.

###### 1.1.5 Summary of experimental conditions

Our experiments provide nine experimental conditions with which to examine the go-or-grow hypothesis:

(1) C8161 in G1, cycling

(2) C8161 in S/G2/M, cycling

(3) C8161 in G1, arrested

(4) WM983C in G1, cycling

(5) WM983C in S/G2/M, cycling

(6) WM983C in G1, arrested

(7) 1205Lu in G1, cycling

(8) 1205Lu in S/G2/M, cycling

(9) 1205Lu in G1, arrested

#### 1.2 Image processing and analysis

Our microscopy data consist of multi-channel time-series stacks of images obtained from 2-D proliferation assays. FUCCI cells in G1 phase appear in the red channel and in S/G2/M phase appear in the green channel. We consider nine experimental conditions: namely, for each of the three cell lines C8161, WM983C and 1205Lu we have cycling cells in G1, cycling cells in S/G2/M, and arrested cells in G1.

It is standard terminology in the microscopy field to refer to an object of interest, such as the image of a cell or cell nucleus, within a microscopy image as a *spot* [16, 17]. Note that the spots in the red and green channels of our microscopy images correspond to the cell nuclei and not to the whole cell. Nevertheless, throughout this document we refer to the red and green spots interchangeably as cell nuclei or cells.

The microscopy data are processed and analysed with Fiji/ImageJ and MATLAB with as much automation as possible. Two main procedures are required. The first procedure is to determine, in each image, the centroids of the spots corresponding to cells in the G1, eS and S/G2/M phases. This requires images which have been processed and segmented to remove as much of the background noise as possible. The second procedure is to obtain 50 cell trajectories for each of the nine experimental conditions using single-cell tracking, and to then use the spot centroids from the first procedure to authenticate the spots identified from tracking. The authentication is used to remove spots in the trajectories which correspond to background noise rather than a cell, and to remove spots from the G1 (red) and S/G2/M (green) trajectories which are actually yellow and therefore appear in both of the red and green channels.

##### 1.2.1 Centroids of cells

The first procedure, which is completely automated within ImageJ and MATLAB, is to identify the spots in the images which correspond to cells and not background noise. This is achieved by preprocessing the images to maximise the signal-to-noise ratio, then thresholding the images to remove the remaining background noise, and finally determining the centroids of all spots in the images. The following describes this in detail.

**Preprocessing:** To maximise the accuracy in identifying spots, which in our case are images of cell nuclei, we enhance the quality of the microscopy images using ImageJ as follows.

1. Import the time-series stack with the Bio-Formats Importer plug-in, splitting the red and green channels.
2. Apply five iterations of Subtract Background with rolling-ball radius of 5 pixels.
3. Apply Enhance Contrast with the Equalize Histogram option selected.
4. Apply the Gaussian Blur filter with sigma = 1.

**Segmentation:** We now identify the spots in the processed images using ImageJ.

1. Apply Auto-thresholding using the Yen method, selecting the option to "calculate the threshold for each image".
2. The resulting binary images are then refined by applying the following commands in the prescribed order.
   (a) Watershed.
   (b) Fill Holes.
   (c) Open, with iterations = 10 and count = 5.
   (d) Watershed.

**Analysis:** For every image in the segmented binary time-series stacks we count the number of spots in each of the red and green channels using ImageJ. We then use MATLAB to determine which spots are yellow.

1. For each of the red and green channels, apply Analyze Particles in ImageJ with size range 5–$\infty$ pixels$^2$ and the option "limit to threshold" selected. Output the stack position and the centroid of every spot in each channel.

2. We now need to determine which spots are red, yellow or green. A spot is red if it appears in the red channel and there is no corresponding spot in the green channel. Similarly, a spot is green if it appears in the green channel and there is no corresponding spot in the red channel. A spot is then yellow if it appears in both of the red and green channels.

   Identifying whether a spot is yellow, and therefore appearing in both of the red and green channels, is complicated by the possible alteration of the shape of the spot during image processing. While we process every image in exactly the same way, the original microscopy images may have different signal-to-noise ratios between the red and green channels. Consequently, there may be a subtle difference in the shape of a spot depending on the channel in which it is viewed, and thereby a difference in the centroid of the spot in each channel.

   We therefore use MATLAB to determine which spots are red, yellow or green, using the stack position and centroid of each spot. The first step is to locate the yellow spots by choosing each red spot in turn and searching the green channel for a corresponding spot in the same stack position as the red spot. Once the yellow spots are identified, the remaining spots are either red or green. Details of this procedure follow below.

   (a) Choose a spot from the red channel and then search the green channel, in the same stack position as the red spot, for a corresponding spot such that the Euclidean distance between the centroids of the two spots is not greater than 3 pixels, noting that the pixel size in our images is 1.8150 $\mu$m. This distance allows for a location error of the centroids of the red and green spots, whereby the centroids may be translated up to one pixel from the original centroid of the yellow spot in the unprocessed images. Placing the original yellow centroid at the centre of a $3 \times 3$ grid of pixels, the red and green centroids from the processed images may be located at any of the nine pixels in the grid. If the green channel has a spot within the specified distance of the red spot then the spot is yellow.

   (b) Once all of the yellow spots are found, the red spots are all of the spots in the red channel which are not yellow. Similarly, the green spots are all of the spots in the green channel which are not yellow.

##### 1.2.2 Single-cell tracking

For each of the nine experimental conditions we require 50 cell trajectories, which we obtain from one randomly-chosen time series, of which there are 30 time-series for cycling cells and 24 time series for G1-arrested cells. We employ single-cell tracking to obtain the trajectories from the red and green channels of our time-series images. The tracking is performed with images which are not thresholded to remove background noise, as thresholding produces binary images in which the spots have lost most of the distinctive features which are required for tracking. Consequently, it is necessary to determine which of the spots identified by the tracking software correspond to cells rather than background noise, and we achieve this by comparing the positions of the tracked spots with the cell centroids that are obtained from thresholded images.

Cell tracking is performed with the TrackMate plug-in for ImageJ [18], using the default settings unless otherwise specified. The tracking is mostly automated within ImageJ, so that the trajectories are identified automatically with TrackMate, however when we select a trajectory for further analysis we manually correct any segmentation or tracking errors. The cell-tracking process is described in detail below.

**Preprocessing:** Within ImageJ, we first prepare the microscopy images for tracking, which includes reducing background noise in the images.

1. Import the time-series stack with the Bio-Formats Importer plug-in, splitting the red and green channels.
2. Apply five iterations of Subtract Background with rolling-ball radius of 5 pixels.
3. For the green channel, apply Enhance Contrast with no options selected, other than to process all slices.
4. For the red channel, modify the Brightness/Contrast settings by applying Reset to restore them to the original values, then apply Set with the "Minimum displayed value" equal to 0 and the "Maximum displayed value" equal to 90. Then select Apply to apply the LUT (lookup table) to all stack slices.

**Tracking:** We use the TrackMate plug-in to find all trajectories in the time-series images, with the following settings.

1. Segmentation is by the Laplacian of Gaussian detection (LoG) algorithm, with the settings detailed in Table S1, and we use neither the median filter nor sub-pixel localisation.

2. We do not filter any spots with initial thresholding of spot quality.

3. For the spot tracker, we use the Linear Assignment Problem (LAP) tracker with the settings detailed in Table S1.

###### 1.2.3 Trajectory selection

Once all of the trajectories in the time series of an experimental condition are identified, we randomly choose trajectories without replacement until 50 trajectories are selected which satisfy prescribed requirements. The requirements for deciding whether to keep or discard a chosen trajectory depend on the particular experimental condition.

**Cycling cells in G1 or S/G2/M:** If the chosen trajectory consists of the complete phase under consideration, therefore either G1 or S/G2/M, then we keep the trajectory. If the trajectory does not consist of the complete phase, such as the trajectory leaves the image area, begins prior to the first frame, or ends after the last frame, then we discard it. If the chosen trajectory has a segmentation or tracking error, and can be manually edited to produce the full trajectory, then we keep it, otherwise we discard it. As our cell-culture conditions ensure minimal G1 arrest, and we randomly choose 50 trajectories from more than $10^3$ trajectories throughout the complete time series, there is little probability of choosing a cell which is in G1 arrest rather than a cycling cell in G1.

**G1-arrested cells:** We need to be able to distinguish between a cell that is arrested in G1 and a cycling cell that is in G1. For each cell line, the time-series images for the G1-arrested cells indicate that almost all of the cells are in G1 phase at a given time, which is very distinct from the cycling cells for which there are cells in all phases of the cell cycle at any given time. It is reasonable to conclude that a randomly-chosen trajectory of a cell from a time series of G1-arrested cells is associated with a G1-arrested cell. Therefore, we keep a chosen trajectory unless it leaves the image area within a duration less than 2 h, as a trajectory of duration less than this is not suitable for our analysis. Most trajectories of G1-arrested cells are much longer than 2 h. If the chosen trajectory has a segmentation or tracking error then we manually edit it.

**Table S1: Settings for the ImageJ plugin TrackMate, for each experimental condition**. The LoG detector settings are for spot segmentation, and the frame-to-frame linking and track segment gap-closing settings are for spot tracking.

| | C8161 G1 | C8161 S/G2/M | C8161 G1 arrest | WM983C G1 | WM983C S/G2/M | WM983C G1 arrest | 1205Lu G1 | 1205Lu S/G2/M | 1205Lu G1 arrest |
|---|---|---|---|---|---|---|---|---|---|
| **LoG detector settings:** | | | | | | | | | |
| Estimated blob diameter ($\mu$m) | 14 | 19 | 22 | 28 | 30 | 26 | 16 | 20 | 26 |
| Threshold | 900 | 300 | 900 | 400 | 200 | 900 | 1200 | 400 | 900 |
| **Frame-to-frame linking:** | | | | | | | | | |
| Maximum distance ($\mu$m) | 50 | 50 | 50 | 50 | 50 | 50 | 50 | 50 | 50 |
| Feature penalties - estimated diameter | 2 | 0 | 2 | 2 | 1 | 2 | 0 | 0 | 2 |
| Feature penalties - total intensity | 2 | 0 | 2 | 2 | 1 | 2 | 0 | 0 | 2 |
| Feature penalties - visibility | 1 | 0 | 1 | 1 | 1 | 1 | 0 | 0 | 1 |
| Feature penalties - quality | 2 | 0 | 2 | 2 | 1 | 2 | 0 | 0 | 2 |
| **Track segment gap-closing:** | | | | | | | | | |
| Maximum distance ($\mu$m) | 50 | 50 | 50 | 50 | 50 | 50 | 50 | 50 | 50 |
| Maximum frame gap | 1 | 1 | 1 | 1 | 1 | 1 | 1 | 1 | 1 |

##### 1.2.4 Trajectory authentication

Each selected trajectory consists of a sequence of spots from either the red or the green channel. Since cells in eS phase appear in both of the red and green channels, and we only want trajectories corresponding to cells in the G1 or S/G2/M phases, we need to remove any spots from the trajectories which are yellow. This is accomplished in MATLAB by comparing the spot positions from tracking with the known centroids of the cells in G1, eS and S/G2/M obtained previously from the thresholded images. Since we are comparing the coordinates of each spot with the centroids of the cells in the same stack position of the time series, it is important to note that while ImageJ usually numbers the frames of a time series starting at one, TrackMate uses frame numbering beginning at zero, so it may be necessary to renumber the frames for each spot of a trajectory to ensure comparison within the same frame.

We set a tolerance of 10 $\mu$m when comparing the spot coordinates with the centroids to allow for the variability associated with estimating positions of spots within images which have been processed differently. Given that the cells have diameters much larger than 10 $\mu$m, for example C8161 cells have a typical diameter of more than 16 $\mu$m [19], the tolerance is relatively small but large enough to find the cell corresponding to a spot.

**G1 trajectories:** Here we detail the authentication of G1 trajectories.

1. Given a trajectory in the red channel, choose each spot in the trajectory in turn.

2. Calculate the Euclidean distance between the coordinates of the spot and the centroids of the cells in the red channel that are in the same stack position as the spot. If there is a cell in eS phase that is within 10 $\mu$m of the spot then we identify the spot as yellow, otherwise the spot corresponds to a cell in G1.

3. Note that any spots identified as yellow will occur towards the end of the trajectory in forward time, corresponding to the cell transitioning from G1 to eS. Therefore, we need to truncate the trajectory so that it corresponds only to the cell in G1 phase. Given that spots corresponding to cells in eS may not always appear yellow due to fluctuations in fluorescence intensity in either channel, we additionally identify a red spot as yellow if it is immediately between two previously identified yellow spots. So, to remove all yellow spots from the trajectory we begin at the final time point of the trajectory and progress backwards in time, removing spots which are yellow

including any spots which are immediately between two previously identified yellow spots. The process terminates when a sequence of two red spots is reached, neither of which are identified as yellow.

**S/G2/M trajectories:** Here we detail the authentication of S/G2/M trajectories.

1. Given a trajectory in the green channel, choose each spot in the trajectory in turn.
2. Calculate the Euclidean distance between the coordinates of the spot and the centroids of the cells in the green channel that are in the same stack position as the spot. If there is a cell in eS phase that is within 10 $\mu$m of the spot then we identify the spot as yellow, otherwise the spot corresponds to a cell in S/G2/M.
3. Note that any spots identified as yellow will occur towards the beginning of the trajectory in forward time, corresponding to the cell transitioning from eS to S/G2/M. Therefore, we need to truncate the trajectory so that it corresponds only to the cell in S/G2/M phase. Given that spots corresponding to cells in eS may not always appear yellow due to fluctuations in fluorescence intensity in either channel, we additionally identify a green spot as yellow if it is immediately between two previously identified yellow spots. So, to remove all yellow spots from the trajectory we begin at the initial time point of the trajectory and progress forwards in time, removing spots which are yellow including any spots which are immediately between two previously identified yellow spots. The process terminates when a sequence of two green spots is reached, neither of which are identified as yellow.

### 2 Data analysis

Here we provide the detailed analysis of the trajectory data for the C8161, WM983C and 1205Lu cell lines.

#### 2.1 Cell-cycle characteristics of each cell line

In Table S2 we provide for each cell line the mean durations of G1 phase, S/G2/M phase, and the total cell cycle for cycling cells [4], and the mean duration of induced G1 arrest (30nM trametinib).

| | C8161 | WM983C | 1205Lu |
|---|---|---|---|
| G1-arrest duration (h) | $34 \pm 15$ | $34 \pm 15$ | $42 \pm 11$ |
| G1 duration (h) | $5 \pm 3$ | $6 \pm 2$ | $18 \pm 9$ |
| S/G2/M duration (h) | $6 \pm 2$ | $8 \pm 3$ | $10 \pm 4$ |
| Total cell-cycle duration (h) | $21 \pm 8$ | $23 \pm 10$ | $37 \pm 16$ |
| Ratio of G1 to S/G2/M durations | 0.9 | 0.7 | 1.8 |
| Ratio of G1 to total durations | 0.3 | 0.2 | 0.5 |
| Ratio of S/G2/M to total durations | 0.3 | 0.4 | 0.3 |

**Table S2: Cell-cycle characteristics for the C8161, WM983C and 1205Lu cell lines**. Data for the cycling cells are from [4], and correspond to 20 cells for each cell line. Data for the G1-arrested cells (30 nM trametinib) are from our current study, and correspond to 50 cells for each cell line. The duration of each cell cycle phase and cell cycle arrest is the mean for the corresponding cell population, and the error is one standard deviation from the mean. The data for the G1-arrest durations correspond to the durations that the cells are arrested during the 48 h of the experiments.

The G1-arrest duration corresponds to the mean duration that the cells are arrested in G1 during the 48 h of the experiment. Note that the cells are likely to be arrested for longer than we can observe during 48 h, so the mean duration that we report corresponds to our observations during the experiment and not necessarily to the total duration for which the cells are arrested. Comparing the long durations of G1 arrest with the relatively short G1 durations of the cycling cells provides quantitative confirmation of the cell-cycle arrest. Indeed, in Figure S1 we show histograms of the durations of G1 arrest for the 50 cells of each cell line.

The data demonstrate that the three cell lines have very different cell-cycle characteristics. In particular, the cell-cycle durations increase in the order C8161, WM983C and 1205Lu. Note that 1205Lu has a relatively long cell cycle, which corresponds to the relatively slow proliferation rate that is observed for this cell line. Further, the ratio of G1 to S/G2/M durations illustrates that C8161 cells spend around the same amount of time in G1 and S/G2/M, WM983C cells spend more time in S/G2/M than G1, and 1205Lu cells spend more time in G1 than S/G2/M, highlighting that these cell lines have very different cell cycles.

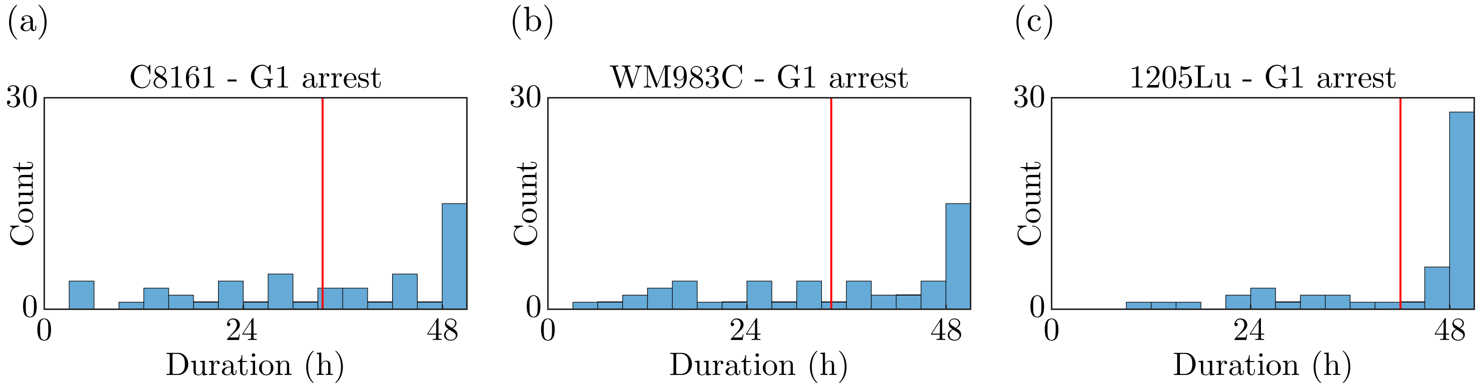

**Figure S1: Histograms of the durations of G1 arrest (30 nM trametinib) for each cell line.** Each histogram corresponds to 50 cells. The vertical red bars indicate the mean durations.

#### 2.2 Cell trajectories

Figure S2 shows trajectories for the WM983C and 1205Lu cell lines, similar to the trajectories for C8161 in Figure 1(g)–(i) in the main document. The insets for Figures S2(c) and (f) show the truncated trajectories

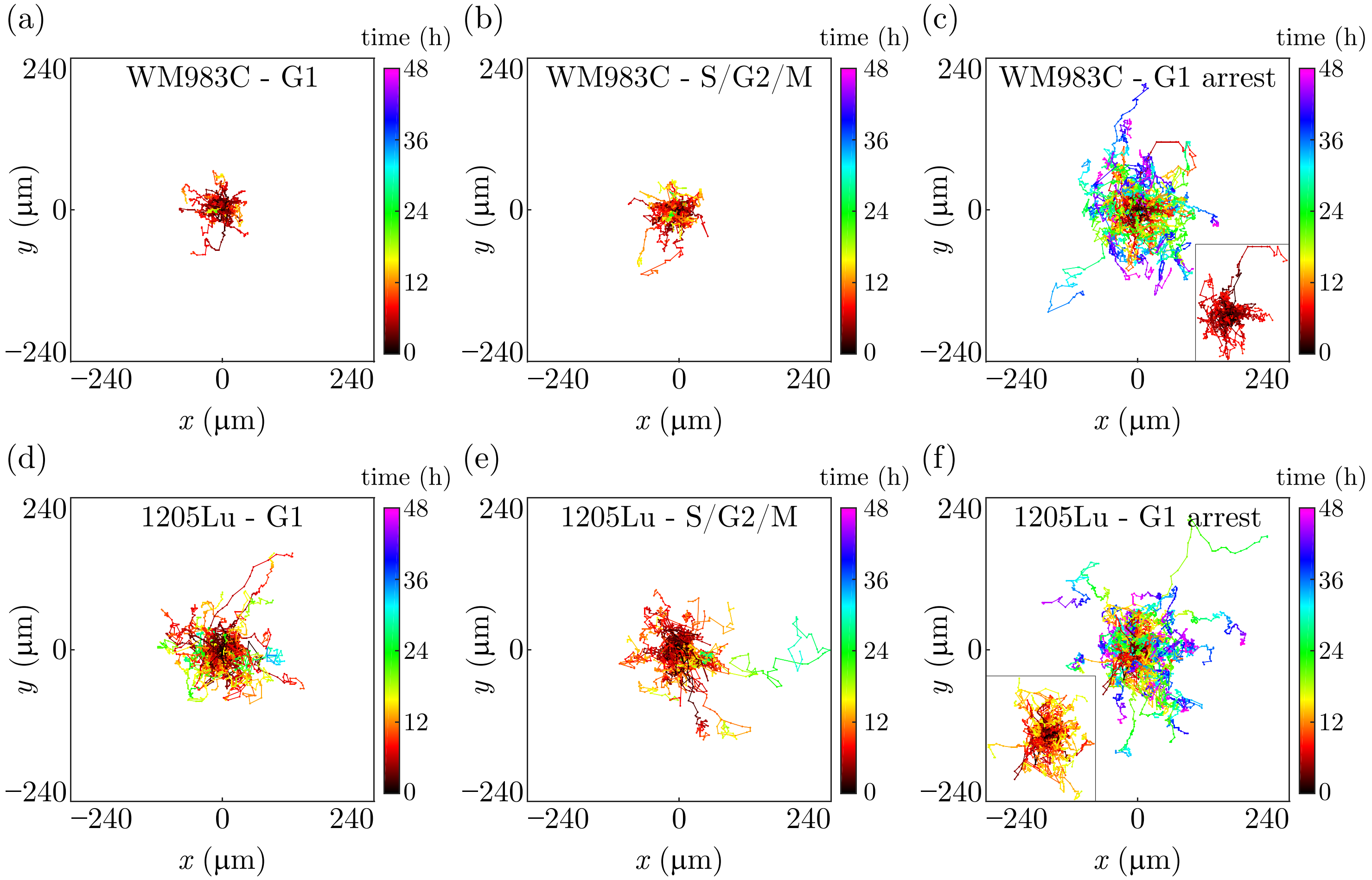

**Figure S2: Cell trajectories of the WM983C and 1205Lu cell lines.** (a)–(c) 50 cell trajectories of G1 cycling, S/G2/M cycling and G1-arrested (30 nM trametinib) WM983C cells, respectively. (d)–(f) 50 cell trajectories of G1 cycling, S/G2/M cycling and G1-arrested (30 nM trametinib) 1205Lu cells, respectively.

for comparison with the cycling cells in G1 given in Figures S2(a) and (d), respectively. The duration of the truncated trajectories is equal to the mean duration of the corresponding trajectories of the cycling cells in G1.

#### 2.3 Cell migration: directionality

We investigate whether the cell migration has directionality by identifying any global anisotropy or directional autocorrelation of the cell trajectories. To reliably identify the presence or absence of anisotropy in cell migration, it is necessary to test for directionality with more than one measure [20,21]. We employ the mean drift velocity and the moment of inertia tensor, which demonstrate that cell migration is globally isotropic for all experimental conditions. We then consider the directional persistence of individual cells over short time intervals, which may not be revealed by measures of global anisotropy, using the temporal velocity autocorrelation function.

##### 2.3.1 Drift velocity

The drift velocity is the mean velocity of the cell from the initial to the final position, and is a measure of the directionality of the trajectory. Specifically, the drift velocity is given by

$$\mathbf{v} = (v_x, v_y), \quad v_x = \frac{x(t_\mathrm{f}) - x(t_\mathrm{i})}{t_\mathrm{f} - t_\mathrm{i}}, \quad v_y = \frac{y(t_\mathrm{f}) - y(t_\mathrm{i})}{t_\mathrm{f} - t_\mathrm{i}}, \tag{S1}$$

where $(x(t), y(t))$ is the cell position at time $t$, $t_\mathrm{i}$ is the time at the initial cell position, and $t_\mathrm{f}$ is the time at the final cell position. By determining the drift velocities in the $x$- and $y$-directions for all trajectories, and then finding the corresponding mean values, we can quantify whether the migrating cells have global directionality. If the mean drift velocity of a cell population is non-zero, then to estimate the diffusivities it is first necessary to subtract the drift from the original cell displacement [22].

Figure S3 shows histograms of the drift velocities, and in Table S3 we statistically characterise the drift velocity data using the mean, median, standard deviation (SD), mean absolute deviation (MAD), the $p$-value of the one-sample $t$-test, the $p$-value of the one-sample sign test, kurtosis, skewness, and the $p$-value of the Anderson-Darling test for normality.

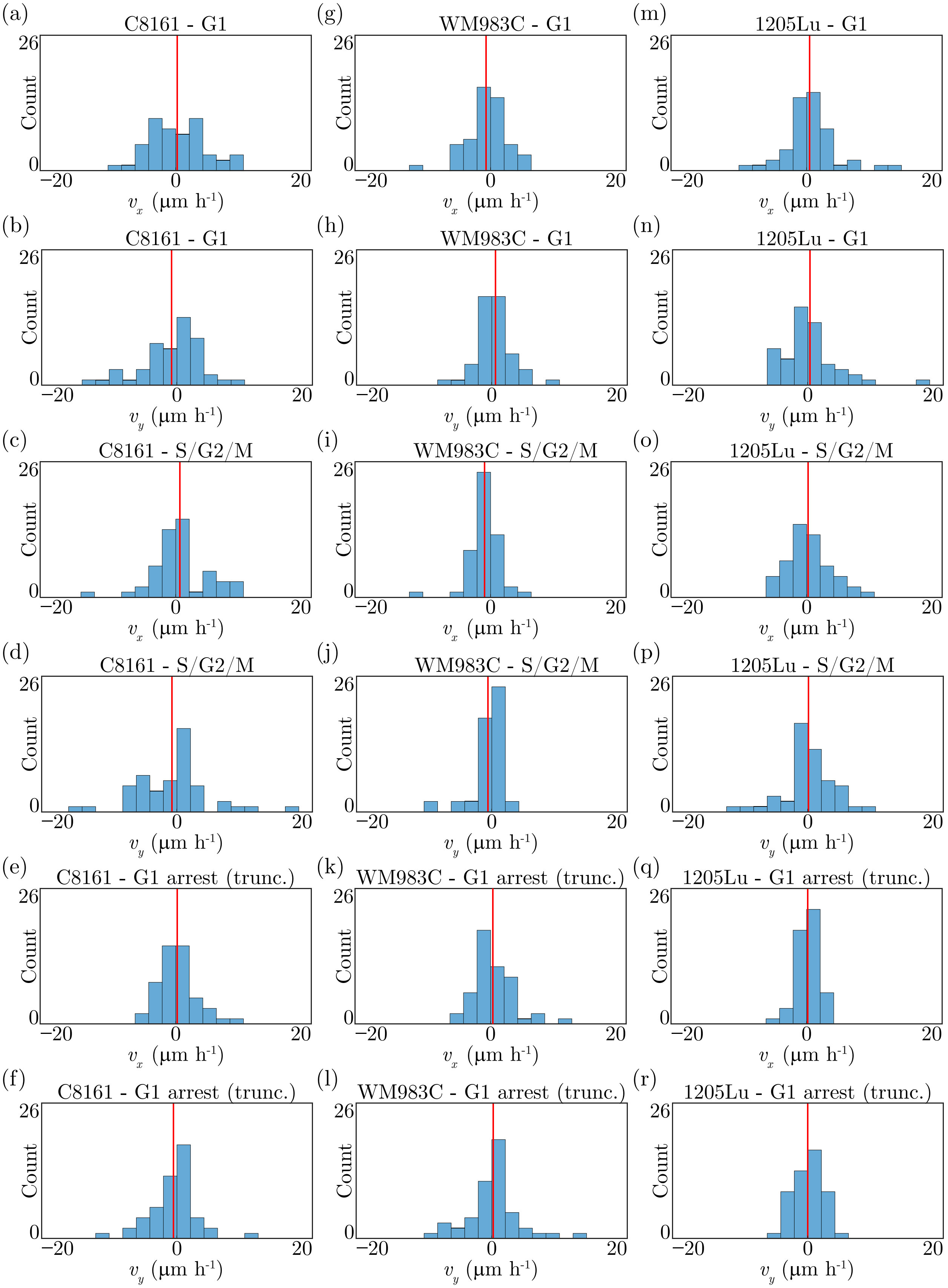

**Figure S3: Drift velocity.** (a) $v_x$ for C8161 - G1. (b) $v_y$ for C8161 - G1. (c) $v_x$ for C8161 - S/G2/M. (d) $v_y$ for C8161 - S/G2/M. (e) $v_x$ for C8161 - G1 arrest (30nM trametinib). (f) $v_y$ for C8161 - G1 arrest (30nM trametinib). (g) $v_x$ for WM983C - G1. (h) $v_y$ for WM983C - G1. (i) $v_x$ for WM983C - S/G2/M. (j) $v_y$ for WM983C - S/G2/M. (k) $v_x$ for WM983C - G1 arrest (30nM trametinib). (l) $v_y$ for WM983C - G1 arrest (30nM trametinib). (m) $v_x$ for 1205Lu - G1. (n) $v_y$ for 1205Lu - G1. (o) $v_x$ for 1205Lu - S/G2/M. (p) $v_y$ for 1205Lu - S/G2/M. (q) $v_x$ for 1205Lu - G1 arrest (30nM trametinib). (r) $v_y$ for 1205Lu - G1 arrest (30nM trametinib). The vertical red bars indicate the mean drift velocity.

**Table S3: Statistical characterisation of the drift-velocity data**.

| | | Mean ($\mu$m h$^{-1}$) | Median ($\mu$m h$^{-1}$) | Standard deviation ($\mu$m h$^{-1}$) | Mean absolute deviation ($\mu$m h$^{-1}$) | One-sample $t$-test $p$-value | One-sample sign test $p$-value | Kurtosis | Skewness | Anderson-Darling test $p$-value |
|---|---|---|---|---|---|---|---|---|---|---|
| **C8161 - G1** | $v_x$ | 0.20 | -0.15 | 4.1 | 3.3 | 1.0 | 0.74 | 3.0 | 0.20 | 0.95 |
| | $v_y$ | -0.79 | 0.33 | 4.3 | 3.3 | 0.20 | 0.89 | 3.6 | -0.73 | 0.0012 |
| **C8161 - S/G2/M** | $v_x$ | 0.59 | 0.16 | 4.2 | 3.0 | 0.33 | 0.89 | 4.4 | -0.083 | 0.019 |
| | $v_y$ | -0.71 | 0.00 | 5.3 | 3.7 | 0.34 | 0.65 | 5.5 | 0.40 | 0.0054 |
| **C8161 - G1 arrest (truncated)** | $v_x$ | 0.24 | -0.16 | 2.8 | 2.1 | 0.55 | 1.0 | 4.7 | 0.81 | 0.30 |
| | $v_y$ | -0.50 | -0.059 | 3.5 | 2.6 | 0.32 | 0.77 | 4.8 | 0.064 | 0.15 |
| **WM983C - G1** | $v_x$ | -0.66 | -0.40 | 2.9 | 2.1 | 0.11 | 0.39 | 4.9 | -0.76 | 0.20 |
| | $v_y$ | 0.52 | 0.54 | 2.5 | 1.8 | 0.15 | 0.67 | 6.0 | 0.23 | 0.058 |
| **WM983C - S/G2/M** | $v_x$ | -0.90 | -0.57 | 2.2 | 1.5 | 0.0060 | 0.0021 | 10 | -1.4 | 0.0014 |
| | $v_y$ | -0.56 | 0.051 | 2.3 | 1.6 | 0.088 | 1.0 | 7.0 | -1.7 | 0.00050 |
| **WM983C - G1 arrest (truncated)** | $v_x$ | 0.33 | -0.21 | 2.9 | 2.2 | 0.43 | 0.67 | 5.2 | 1.1 | 0.023 |
| | $v_y$ | 0.21 | 0.32 | 3.9 | 2.6 | 0.71 | 0.47 | 5.3 | 0.51 | 0.0027 |
| **1205Lu - G1** | $v_x$ | 0.43 | 0.21 | 3.8 | 2.7 | 0.43 | 0.57 | 5.5 | 0.81 | 0.0091 |
| | $v_y$ | 0.33 | -0.068 | 4.2 | 2.9 | 0.58 | 0.67 | 7.0 | 1.5 | 0.011 |
| **1205Lu - S/G2/M** | $v_x$ | 0.24 | -0.068 | 3.3 | 2.7 | 0.61 | 1.0 | 2.9 | 0.35 | 0.49 |
| | $v_y$ | 0.084 | 0.057 | 3.7 | 2.6 | 0.87 | 1.0 | 4.9 | -0.60 | 0.024 |
| **1205Lu - G1 arrest (truncated)** | $v_x$ | 0.13 | 0.27 | 1.7 | 1.4 | 0.61 | 0.48 | 3.9 | -0.62 | 0.79 |
| | $v_y$ | -0.017 | 0.055 | 2.3 | 1.8 | 0.96 | 0.78 | 2.7 | -0.15 | 0.61 |

We use the one-sample $t$-test to determine whether the mean of each drift-velocity component differs statistically from a population with mean of 0 $\mu$m h$^{-1}$. With a cut-off of 0.05 for statistical significance, the $p$-values indicate that, except in the case of $v_x$ for WM983C in S/G2/M, the results are not statistically significant. Regarding $v_x$ for WM983C in S/G2/M, we find that the results are not statistically significant with a cut-off of 0.05 when we hypothesise a population mean of -0.3 $\mu$m h$^{-1}$. Over the complete 48 h of the experiments, a drift of magnitude 0.3 $\mu$m h$^{-1}$ corresponds to a displacement of magnitude 14.4 $\mu$m, which is less than the diameter of a typical melanoma cell, such as 16.44 $\mu$m for C8161 [19]. Therefore, we conclude that there is no drift in any of our migration experiments. We arrive at the same conclusion with a similar analysis using the one-sample sign test and the median of each drift-velocity component.

Kurtosis is a measure of how light-tailed or heavy-tailed a distribution is relative to the normal distribution. The value of the kurtosis for a normal distribution is 3, which indicates that our drift-velocity data are generally heavy-tailed.

Skewness is a measure of the asymmetry of a distribution, so that a symmetric distribution such as the normal distribution has a skewness of zero. Our drift-velocity data generally have skewness values that differ from zero, indicating that the data have a degree of asymmetry.

The Anderson-Darling test provides a measure for whether each set of drift-velocity data is a sample from a normal distribution. Assuming that the sample data are normally distributed, the calculated $p$-value is the conditional probability of observing data at least as extreme as the sample data under consideration. Therefore, the $p$-value is a measure of the strength of evidence against the assumption that the sample data are from a normal distribution. Applying the test to our data, with a cut-off of 0.05 for statistical significance, we conclude that around half of the data sets are unlikely to be sampled from a normal distribution. We therefore cannot assume in general that the drift-velocity data are distributed normally.

##### 2.3.2 Moment of inertia tensor

The moment of inertia tensor $\mathbf{I}$ can be used to reliably identify anisotropy in cell migration [20, 21, 23]. Here we consider the mean moment of inertia tensor $\bar{\mathbf{I}}$, which quantifies the anisotropy of the mean of all cell trajectories.

For the 50 trajectories of each experimental condition we consider all of the displacements that occur during the 15 min time interval between images in the time series. We denote these displacements by $(x_i, y_i)$, for $i = 1, \ldots, N$, where $N$ is the total number of displacements. With the start point at the origin, we consider a unit mass at the end point of each displacement. The two-dimensional mean moment of inertia tensor is then given by

$$\bar{\mathbf{I}} = \begin{pmatrix} I_{xx} & I_{xy} \\ I_{yx} & I_{yy} \end{pmatrix}, \quad \text{where} \quad I_{xx} = \sum_{i=1}^{N} y_i^2, \quad I_{xy} = I_{yx} = -\sum_{i=1}^{N} x_i y_i \quad \text{and} \quad I_{yy} = \sum_{i=1}^{N} x_i^2. \tag{S2}$$

The eigenvalues $\overline{\lambda}_1$ and $\overline{\lambda}_2$ of $\bar{\mathbf{I}}$, which are the principal moments of inertia, indicate directionality in cell migration. By convention $\overline{\lambda}_1 > \overline{\lambda}_2$, so that $\overline{\lambda}_1 = \overline{\lambda}_2$ corresponds to isotropy, and $\overline{\lambda}_1 > \overline{\lambda}_2$ corresponds to anisotropy. The ratio $\overline{\lambda}_1/\overline{\lambda}_2 \geq 1$ therefore quantifies the degree of anisotropy, or directionality, in cell migration. Table S4 provides the ratio $\overline{\lambda}_1/\overline{\lambda}_2$ for cycling cells in G1 and S/G2/M, and for cells arrested in G1 with trametinib (30 nM), for each of the three cell lines. The results indicate that $\overline{\lambda}_1/\overline{\lambda}_2$ is almost equal to one for all experimental conditions, hence we conclude that the cell migration is essentially globally isotropic for all conditions. This result is consistent with the finding of no drift in all experiments. We also include the results for the G1-arrested cells over the full 48 h of the experiments, which show that there is no anisotropy over the complete experiment duration.

The error estimates in Table S4 are obtained by bootstrapping the set of all $x$-displacements and the set of all $y$-displacements, each with $10^4$ iterations. Each re-sampling of the displacements is used to calculate $\bar{\mathbf{I}}$ and $\overline{\lambda}_1/\overline{\lambda}_2$. The standard deviation of the $10^4$ values of $\overline{\lambda}_1/\overline{\lambda}_2$ is then taken to be the error.

| | $\overline{\lambda}_1/\overline{\lambda}_2$ |
|---|---|
| C8161 - G1 | $1.18 \pm 0.09$ |
| C8161 - S/G2/M | $1.39 \pm 0.12$ |
| C8161 - G1 arrest | $1.10 \pm 0.05$ |
| C8161 - G1 arrest (truncated) | $1.03 \pm 0.07$ |
| WM983C - G1 | $1.15 \pm 0.07$ |
| WM983C - S/G2/M | $1.06 \pm 0.08$ |
| WM983C - G1 arrest | $1.22 \pm 0.06$ |
| WM983C - G1 arrest (truncated) | $1.29 \pm 0.10$ |
| 1205Lu - G1 | $1.04 \pm 0.04$ |
| 1205Lu - S/G2/M | $1.32 \pm 0.07$ |
| 1205Lu - G1 arrest | $1.03 \pm 0.03$ |
| 1205Lu - G1 arrest (truncated) | $1.06 \pm 0.06$ |

**Table S4: Ratio of the eigenvalues for the mean moment of inertia tensor**. The error estimates are obtained by bootstrapping the sets of $x$- and $y$-displacements, each with $10^4$ iterations, and calculating $\overline{\lambda}_1/\overline{\lambda}_2$ for each re-sampled data set, whereby the error is then taken as the standard deviation of the $10^4$ values of $\overline{\lambda}_1/\overline{\lambda}_2$.

##### 2.3.3 Temporal velocity autocorrelation function

The temporal velocity autocorrelation function (TVAF) provides a measure for directional persistence of cell migration [24–26]. We say that the velocity $\mathbf{v}(t)$ of a cell at time $t$ is equal to the mean velocity of the cell between the time points $t$ and $t + \Delta t$, where $t$ is a time point of the experimental time-series images and $\Delta t$ is the time interval between images, here 15 min. If $\mathbf{r}(t)$ is the position of the cell at time $t$, then we have $\mathbf{v}(t) = \big(\mathbf{r}(t + \Delta t) - \mathbf{r}(t)\big)/\Delta t$. The TVAF is given by

$$C_v(\tau) = \big\langle \mathbf{v}(t + \tau) \cdot \mathbf{v}(t) \big\rangle, \tag{S3}$$

where, for a given lag time $\tau$, the averaging is over all possible time points $t$ of a cell trajectory and over all cell trajectories. The normalised, and dimensionless, TVAF is

$$\widehat{C}_v(\tau) = \frac{C_v(\tau)}{C_v(0)} = \frac{\big\langle \mathbf{v}(t + \tau) \cdot \mathbf{v}(t) \big\rangle}{\big\langle \mathbf{v}(t) \cdot \mathbf{v}(t) \big\rangle}. \tag{S4}$$

For a population of cells in which the migration can be described as a random walk, $\widehat{C}_v$ will be 1 at $\tau = 0$ and 0 for all $\tau > 0$, revealing no correlation in cell migration. A persistent random walk is revealed by a continuously decaying profile for $\widehat{C}_v$, since the correlation reduces over time. To estimate the decorrelation time for the persistence, we fit to $\widehat{C}_v$ either a single exponential function, $f_1$, or a sum of two exponential functions, $f_2$, which we write generally as

$$f_n(\tau) = \sum_{i=1}^{n} A_i e^{-\tau/\mathcal{T}_i} \text{ , for } n = 1 \text{ or } 2, \tag{S5}$$

and constants $A_i$ and $\mathcal{T}_i$, where the latter are the decorrelation times. When $n = 2$ we adopt the convention that $\mathcal{T}_1 < \mathcal{T}_2$. We estimate the decorrelation times $\mathcal{T}_i$ by fitting Equation (S5) to $\widehat{C}_v$ in Equation (S4) evaluated with experimental data, using the `fit` function and either the `exp1` or the `exp2` models [27] in MATLAB. A best fit is obtained with $n = 2$ for the cell lines C8161 and 1205Lu, and $n = 1$ for the WM983C cell line, shown in Figure S4 and Table S5. Therefore, for the cell lines C8161 and 1205Lu, a sum of two exponential

| | $\mathcal{T}_1$ (h) | $\mathcal{T}_2$ (h) |
|---|---|---|
| C8161 - G1 | $0.10 \pm 0.03$ | $4 \pm 1$ |
| C8161 - S/G2/M | $0.18 \pm 0.02$ | $3 \pm 3$ |
| C8161 - G1 arrest (truncated) | $0.010 \pm 0.003$ | $4.0 \pm 0.6$ |
| WM983C - G1 | $0.11 \pm 0.01$ | - |
| WM983C - S/G2/M | $0.010 \pm 0.001$ | - |
| WM983C - G1 arrest (truncated) | $0.09 \pm 0.01$ | - |
| 1205Lu - G1 | $0.13 \pm 0.01$ | $2.2 \pm 0.7$ |
| 1205Lu - S/G2/M | $0.003 \pm 0.003$ | $0.15 \pm 0.08$ |
| 1205Lu - G1 arrest (truncated) | $0.006 \pm 0.003$ | $0.15 \pm 0.02$ |

**Table S5: Estimated decorrelation times of directional persistence.** The temporal velocity autocorrelation function data is fit with a sum of two exponential functions for C8161 and 1205Lu, and is fit with a single exponential function for WM983C. The error estimates are obtained by a bootstrapping process which produces $10^4$ estimates of $\widehat{C}_v$, from which decorrelation times are estimated by fitting Equation (S5). The error is then taken to be the standard deviation of the $10^4$ estimates for the decorrelation times.

functions is found to fit the TVAF data best, whereas for the WM983C cell line, a best fit is obtained with a single exponential function.

Both the exponential model and the sum of two exponentials model have previously been observed as the functional forms for the TVAF in adherent cell migration on two-dimensional surfaces [26, 28]. Our results suggest that the WM983C cells exhibit a single slow decorrelation time, while the C8161 and 1205Lu cells exhibit both a slow and a fast decorrelation time. For the 1205Lu cell line, the G1-arrested cells appear to decorrelate faster than the cycling cells in G1. These observations illustrate that similar cell types, in this case melanoma cells, can have distinctly different migratory behaviour both with and without drug-induced

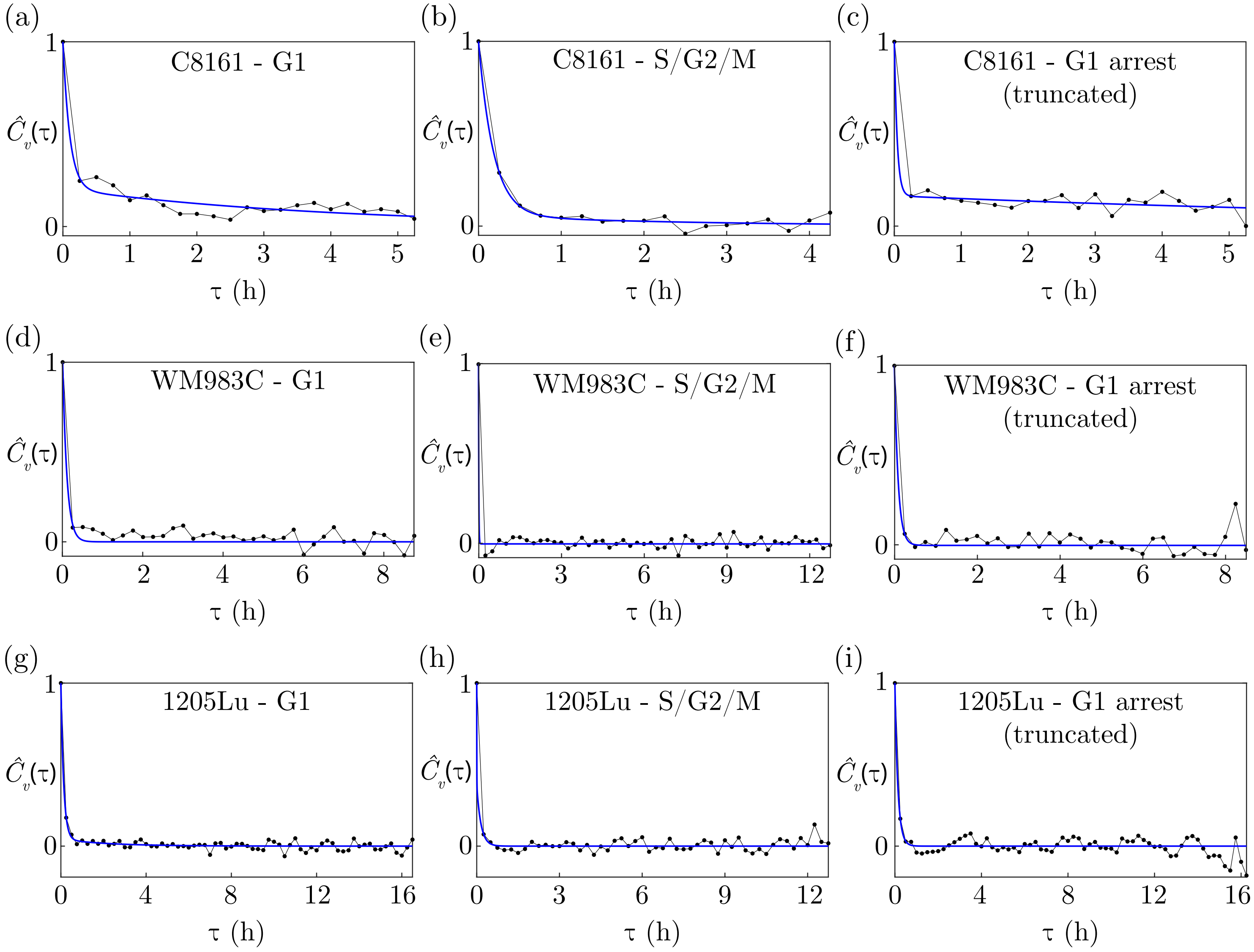

**Figure S4: Best fit of the exponential function, $f_1$, or the sum of two exponential functions, $f_2$, in Equation (S5) to the normalised temporal velocity autocorrelation function, $\widehat{C}_v(\tau)$, in Equation (S4) evaluated with experimental data.** (a)–(c) G1 cycling, S/G2/M cycling, and truncated G1-arrested (30nM trametinib) C8161 cells, respectively, for which $f_2$ fits best. (d)–(f) G1 cycling, S/G2/M cycling, and truncated G1-arrested (30nM trametinib) WM983C cells, respectively, for which $f_1$ fits best. (g)–(i) G1 cycling, S/G2/M cycling, and truncated G1-arrested (30nM trametinib) 1205Lu cells, respectively, for which $f_2$ fits best. The black discs correspond to the data and the blue curves to the best fit of $f_1$ or $f_2$.

cell cycle arrest.

The estimated error in the decorrelation times, indicated in Table S5, is obtained using a bootstrapping process. For each lag time $\tau$, we re-sample the data set $\{\, \mathbf{v}(t+\tau)\cdot\mathbf{v}(t) \mid \text{for all possible } t \,\}$ with replacement, to give a re-sampled data set of the same size. Using the mean values of each of these re-sampled data sets, we calculate $\widehat{C}_v$ and then fit Equation (S5), thereby obtaining new estimates for the decorrelation times. The re-sampling process is performed $10^4$ times to obtain $10^4$ estimates for the decorrelation times, and the standard deviation of these estimates is taken to be the error in the original estimates of the decorrelation times.

Fitting a function which is a sum of two exponentials to data is well known to be ill-conditioned [29]. Consequently, the estimated errors in the decorrelation times may be proportionally larger when using a model which is the sum of two exponentials compared with a single exponential model. This has minimal consequence for our purposes as we only need an estimate of the decorrelation times as a guide for the smallest lag time after which the cell migration is essentially random, allowing us to then estimate diffusivities.

Based on the decorrelation times in Table S5, we note that most of the correlation is lost within 1 h, for all cell lines. While 1205Lu and particularly C8161 each have an associated slow decorrelation time which results in residual correlation beyond 1 h, it is not practical to use longer lag times for estimating the diffusivities due to the limitations imposed by the durations of the G1 and S/G2/M phases. Any residual correlation beyond a lag time of 1 h, however, is relatively small and would have minimal consequence for our estimation of diffusivities.

#### 2.4 Cell migration: diffusivities

The migration of an adherent cell tends to exhibit directional persistence, as we demonstrate with the temporal velocity autocorrelation function (TVAF) applied to our three melanoma cell lines. Cell migration is therefore often modelled as a persistent random walk (PRW) [22, 30, 31]. There are, however, numerous experimental studies that do not support the PRW as a model for cell migration, as the characteristics of the observed migration differ from predictions of the PRW models [26, 28, 32, 33]. In particular, PRW models predict that the TVAF has a simple exponential form [26,28]. For our cell lines, however, only one cell line has a TVAF with an exponential form, while the other two cell lines have a TVAF with the form of a sum of two exponentials. Therefore, we do not employ a PRW model to estimate diffusivities. Rather, our diffusivity estimates are obtained over time periods for which the directional persistence is lost and the cells are undergoing free diffusion.

We estimate the diffusivities using the mean square displacement (MSD), which has various definitions [34–36]. Here we discuss our methodology in detail. Consider a cell trajectory in 2-D obtained from a time series of $N$ images, where $\Delta t$ is the time interval between successive positions. In our case, $\Delta t = 0.25$ h. We denote the sequence of cell positions as $\{\, \mathbf{r}_i \,\}_{i=1}^{N}$. Let $1 \leq M \leq N - 1$ be the integer such that $M\Delta t$ is the specified minimum lag time. For the lag time $\tau_n = n\Delta t$, where $n = M, \ldots, N - 1$, there are $N - n$ non-zero forward displacements along the cell trajectory. The MSD, denoted $\rho(\tau_n)$, is given by

$$\rho(\tau_n) = \frac{1}{N-n} \sum_{i=1}^{N-n} \|\mathbf{r}_{i+n} - \mathbf{r}_i\|^2. \tag{S6}$$

We have established that our cell migration experiments are globally isotropic, and that the directional persistence of the cells dissipates within 1 h. Therefore, we may regard the cell migration as free diffusion for lag times of at least 1 h, which corresponds to $M = 4$. We may then estimate the diffusivity $D$ of the cell by fitting the equation [35]

$$\rho(\tau) = 4D\tau \text{ , for lag time } \tau, \tag{S7}$$

to the set of data points $\{ \big(\tau_n, \rho(\tau_n)\big) \mid n = M, \ldots, N-1 \}$. We fit Equation (S7) to our data using the `fitlm` linear regression function in MATLAB [37] with no constant term, therefore passing through the origin.

For each cell line and experimental condition we have 50 cell trajectories. For each trajectory we calculate $D$, using the minimum lag time of 1 h, within the 2-h time intervals 0–2 h, 1–3 h, 2–4 h and so on, successively offset by 1 h up until the end of the trajectory. To illustrate the variability in $D$ for the 50 cell trajectories for each cell line and experimental condition we show in Figure S5 the histograms of $D$ within the time interval 0–2 h. For each 2-h time interval we then calculate the mean diffusivity $\langle D \rangle$ from the individual $D$ corresponding to all trajectories that extend to the end of the interval. Note that the mean may be taken over fewer than 50 trajectories due to the differing durations of the trajectories. Figure 1(j)–(l) in the main document shows a comparison of $\langle D \rangle$ as a function of time interval between cycling cells in G1, cycling cells in S/G2/M, and G1-arrested cells (30 nM trametinib), for each of the cell lines C8161, WM983C and 1205Lu.

For each cell line we have three independent samples of diffusivities corresponding to three experimental conditions: cycling cells in G1, cycling cells in S/G2/M, and G1-arrested cells (30 nM trametinib). To compare the mean diffusivities between each pair of diffusivity samples we employ the permutation test, which is a non-parametric significance test which makes no assumptions about the population distributions such as normality and equal variance. For each cell line we are interested in three null hypotheses, $H_0$, where we denote the mean diffusivities of cycling cells in G1 as $\langle D \rangle_{\text{G1}}$, of cycling cells in S/G2/M as $\langle D \rangle_{\text{S/G2/M}}$, and of G1-arrested cells as $\langle D \rangle_{\text{G1-arrest}}$:

1. $H_0 : \langle D \rangle_{\text{G1}} = \langle D \rangle_{\text{S/G2/M}}$, that is, the mean diffusivities are the same for cells in G1 and cells in S/G2/M.

   Since we are interested in whether the mean diffusivities are equal, we test if $\langle D \rangle_{\text{G1}}$ is significantly greater

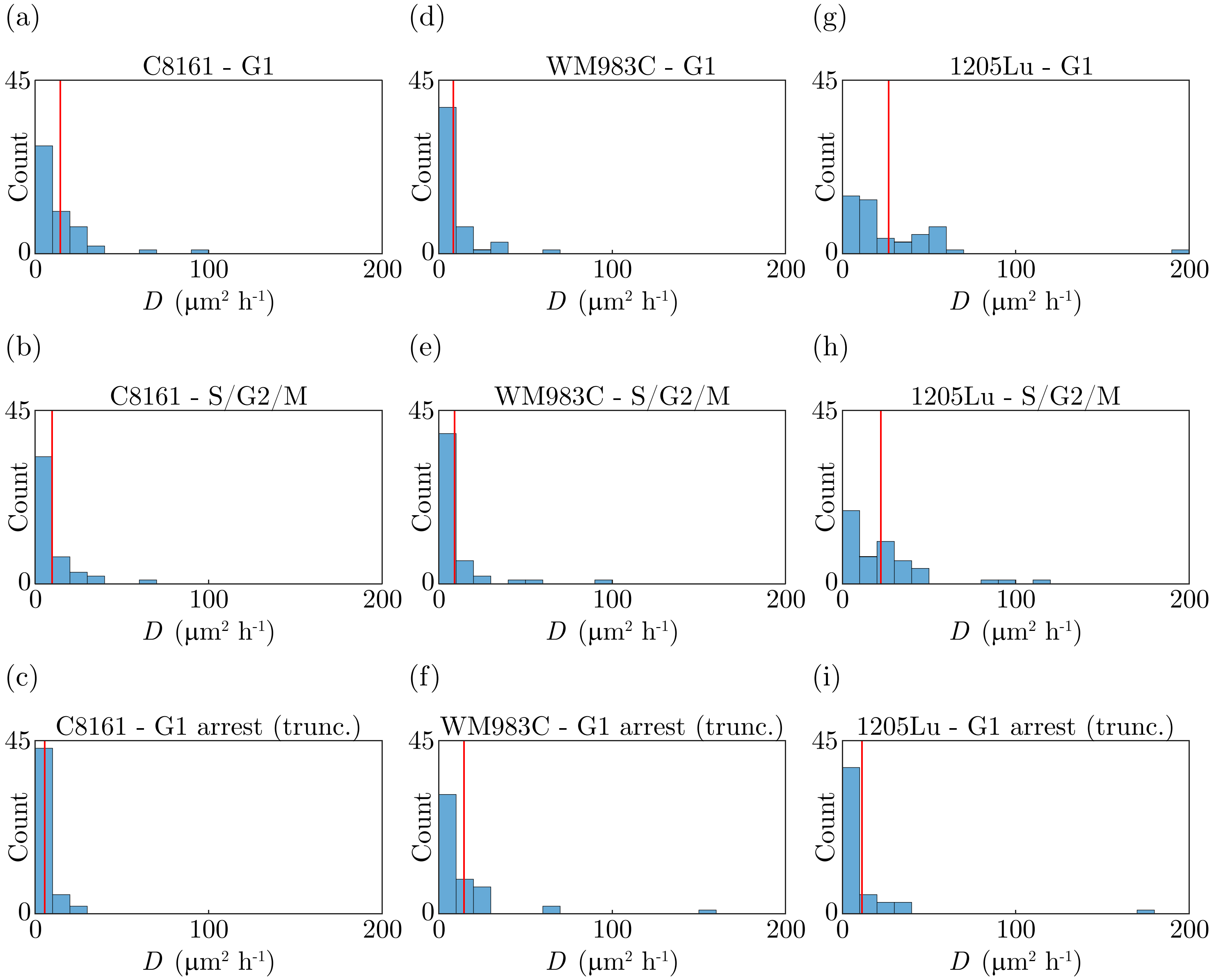

**Figure S5: Histograms of the diffusivities $D$ within the time interval 0–2 h for each cell line and experimental condition.** Each histogram corresponds to 50 cell trajectories. The vertical red bars indicate $\langle D \rangle$.

than $\langle D \rangle_{\mathrm{S/G2/M}}$ and if $\langle D \rangle_{\mathrm{S/G2/M}}$ is significantly greater than $\langle D \rangle_{\mathrm{G1}}$, which is therefore a two-sided test.

2. $H_0 : \langle D \rangle_{\mathrm{G1}} \geq \langle D \rangle_{\mathrm{G1\text{-}arrest}}$, that is, the mean diffusivity for G1-arrested cells is not larger than the mean diffusivity for cycling cells in G1. Since the G1-arrested cells are not progressing through the cell cycle, the go-or-grow hypothesis suggests that they may be more motile, with correspondingly larger mean diffusivities, than the cycling cells. So we test whether $\langle D \rangle_{\mathrm{G1\text{-}arrest}}$ is significantly greater than $\langle D \rangle_{\mathrm{G1}}$, which is therefore a one-sided test.

3. $H_0 : \langle D \rangle_{\mathrm{S/G2/M}} \geq \langle D \rangle_{\mathrm{G1\text{-}arrest}}$, that is, the mean diffusivity for G1-arrested cells is not larger than the mean diffusivity for cycling cells in S/G2/M. Since the G1-arrested cells are not progressing through

the cell cycle, the go-or-grow hypothesis suggests that they may be more motile, with correspondingly larger mean diffusivities, than the cycling cells. So we test whether $\langle D \rangle_{\text{G1-arrest}}$ is significantly greater than $\langle D \rangle_{\text{S/G2/M}}$, which is therefore a one-sided test.

In Table S6 we provide the $p$-values, with the corresponding effect sizes in parentheses, obtained from the permutation test for each time interval. The effect size is defined as the magnitude of the difference of the two sample means divided by the mean of the two sample standard deviations. If we use a threshold of 0.01 for statistical significance, we note that in almost all cases the $p$-values indicate a lack of significance, so we cannot reject the corresponding null hypotheses. These tests don't prove that the null hypotheses are true, however we are unable to reject them based on our current data. For WM983C the hypothesis tests indicate that $\langle D \rangle_{\text{G1-arrest}}$ is significantly greater than $\langle D \rangle_{\text{G1}}$ for two of the eight time intervals, and $\langle D \rangle_{\text{G1-arrest}}$ is significantly greater than $\langle D \rangle_{\text{S/G2/M}}$ for three of the eight time intervals. The particular threshold of 0.01 for statistical significance is arbitrary, and there is increasing support to employ a threshold of 0.005 for new scientific discoveries in order to reduce false positives and improve the reproducibility of research [38]. If we employ a threshold of 0.005 for statistical significance then the hypothesis tests indicate that, for WM983C, $\langle D \rangle_{\text{G1-arrest}}$ is not significantly greater than $\langle D \rangle_{\text{G1}}$ for any time interval, and $\langle D \rangle_{\text{G1-arrest}}$ is significantly greater than $\langle D \rangle_{\text{S/G2/M}}$ for only two of the eight time intervals. While statistical significance can be a helpful guide, the ultimate interest in biology must be the biological importance of an effect [39], on which significance testing provides no information. In our case, the significance testing indicates an effect in only a relatively few time intervals for WM983C. The data in Figure 1(j)–(l) and Figure S5 demonstrate that the mean diffusivities for each cell line are remarkably similar between the different experimental conditions, where typically one mean diffusivity is within one standard deviation of another mean diffusivity. Therefore, there appears to be no biological difference between the mean diffusivities of the three experimental conditions.

| Time interval (h) | 0–2 | 1–3 | 2–4 | 3–5 | 4–6 | 5–7 | 6–8 | 7–9 |
|---|---|---|---|---|---|---|---|---|
| **C8161** | | | | | | | | |
| $H_0 : \langle D \rangle_{\text{G1}} = \langle D \rangle_{\text{S/G2/M}}$ | 0.11 (0.34) | 0.57 (0.12) | 0.63 (0.12) | 0.44 (0.26) | 0.89 (0.077) | - | - | - |
| $H_0 : \langle D \rangle_{\text{G1}} \geq \langle D \rangle_{\text{G1-arrest}}$ | 1.0 (0.80) | 1.0 (0.83) | 1.0 (0.64) | 1.0 (0.64) | 0.76 (0.22) | - | - | - |
| $H_0 : \langle D \rangle_{\text{S/G2/M}} \geq \langle D \rangle_{\text{G1-arrest}}$ | 0.99 (0.47) | 1.0 (0.64) | 1.0 (0.72) | 0.95 (0.46) | 0.77 (0.26) | - | - | - |
| **WM983C** | | | | | | | | |
| $H_0 : \langle D \rangle_{\text{G1}} = \langle D \rangle_{\text{S/G2/M}}$ | 0.77 (0.063) | 0.92 (0.021) | 0.49 (0.14) | 0.18 (0.29) | 0.86 (0.039) | 0.68 (0.099) | 0.094 (0.39) | 0.50 (0.18) |
| $H_0 : \langle D \rangle_{\text{G1}} \geq \langle D \rangle_{\text{G1-arrest}}$ | 0.046 (0.35) | 0.11 (0.27) | 0.055 (0.35) | 0.12 (0.25) | 0.0069 (0.56) | 0.021 (0.50) | 0.40 (0.080) | 0.0065 (0.67) |
| $H_0 : \langle D \rangle_{\text{S/G2/M}} \geq \langle D \rangle_{\text{G1-arrest}}$ | 0.11 (0.27) | 0.099 (0.29) | 0.012 (0.48) | 0.0035 (0.59) | 0.0054 (0.54) | 0.025 (0.41) | 0.038 (0.41) | 0.0046 (0.58) |
| **1205Lu** | | | | | | | | |
| $H_0 : \langle D \rangle_{\text{G1}} = \langle D \rangle_{\text{S/G2/M}}$ | 0.42 (0.17) | 0.94 (0.016) | 0.83 (0.043) | 0.59 (0.13) | 0.27 (0.25) | 0.50 (0.19) | 0.92 (0.023) | 0.34 (0.22) |
| $H_0 : \langle D \rangle_{\text{G1}} \geq \langle D \rangle_{\text{G1-arrest}}$ | 1.0 (0.55) | 1.0 (0.58) | 0.90 (0.28) | 0.80 (0.21) | 0.88 (0.26) | 0.97 (0.37) | 1.0 (0.69) | 1.0 (0.85) |
| $H_0 : \langle D \rangle_{\text{S/G2/M}} \geq \langle D \rangle_{\text{G1-arrest}}$ | 0.99 (0.44) | 1.0 (0.57) | 0.91 (0.28) | 0.86 (0.25) | 0.62 (0.11) | 0.90 (0.29) | 1.0 (0.70) | 1.0 (0.9) |

**Table S6: Permutation tests on mean diffusivities $\langle D \rangle$ for the C8161, WM983C and 1205Lu cell lines.** For each experimental condition, cycling G1, cycling S/G2/M and G1 arrest, and relevant time interval, we provide the $p$-value and corresponding effect size (in parentheses) obtained from the permutation test.

We conclude that:

- For each cell line and experimental condition, $\langle D \rangle$ has little variation over time, hence motility is essentially constant during each cell-cycle phase and during G1 arrest.

- For each cell line there is remarkably little variation in $\langle D \rangle$ between cycling cells in G1, cycling cells in S/G2/M, and G1-arrested cells. Therefore, cell motility is the same whether the cells are in G1, S/G2/M or G1-arrested.

- The three cell lines have very different proliferation and migration characteristics, however $\langle D \rangle$ is remarkably consistent across the cell lines.

In Figure S6 we show $\langle D \rangle$ as a function of time interval over the full 48 h of the experiments for all three cell lines in G1-arrest (30 nM trametinib). The data in Figure S6 demonstrate the remarkably low variability of $\langle D \rangle$ over the experimental duration of 48 h. For each cell line, the cells are arrested in G1 and so are not in a proliferative state, however we observe essentially no change in motility over two days.

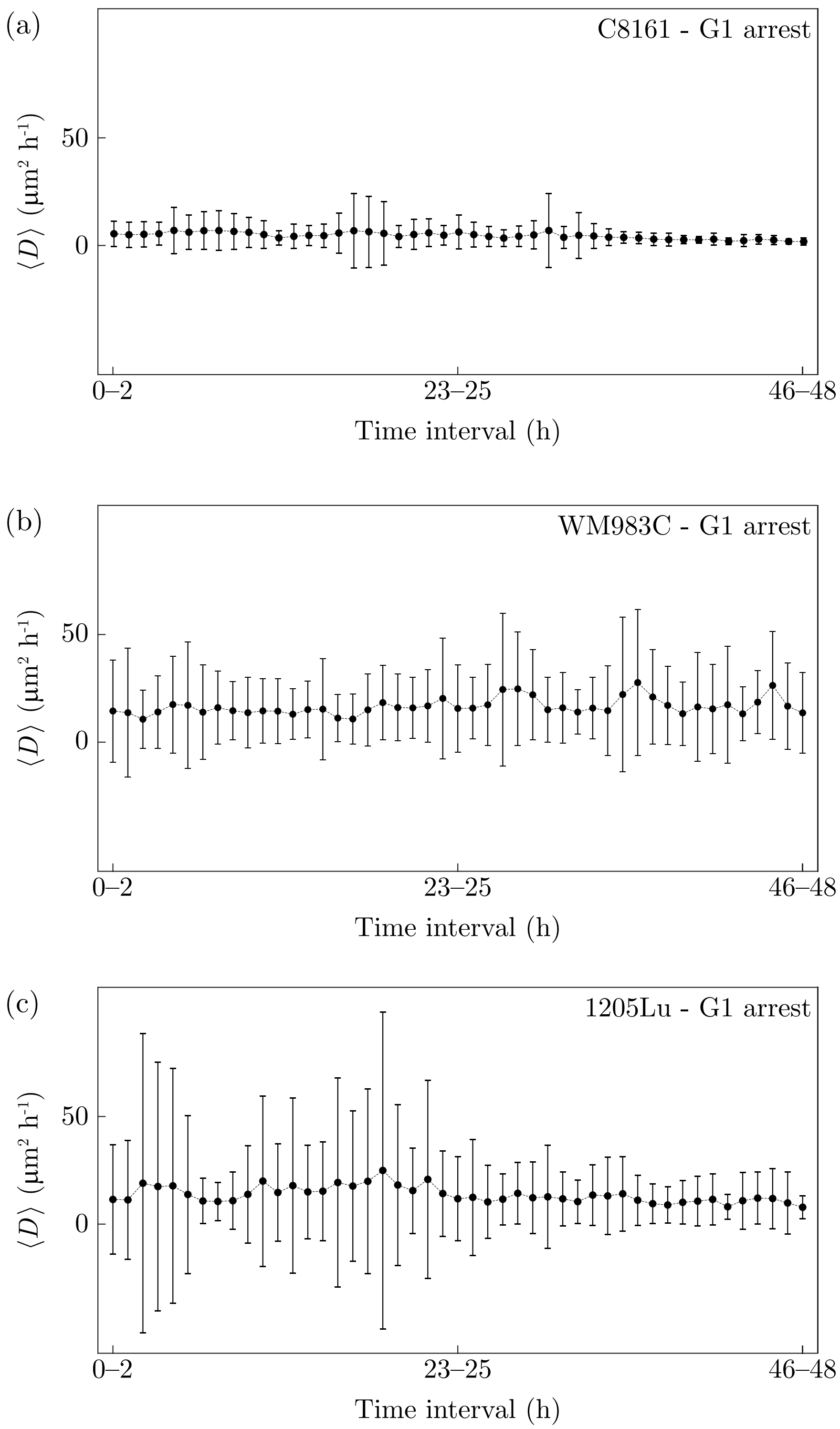

**Figure S6: Mean diffusivities for C8161, WM983C and 1205Lu in G1 arrest (30nM trametinib) over 48 h.** (a)–(c) $\langle D \rangle$ for C8161, WM983C and 1205Lu, respectively, for the range of time intervals over the full experiment duration of 48 h. In each case we show $\langle D \rangle$, and report the variability using $\langle D \rangle$ plus or minus the sample standard deviation.
